# Lack of Cas13a inhibition by anti-CRISPR proteins from *Leptotrichia* prophages

**DOI:** 10.1101/2021.05.27.445852

**Authors:** Alexander J Meeske, Matthew C. Johnson, Logan T. Hille, Benjamin P. Kleinstiver, Joseph Bondy-Denomy

**Affiliations:** Department of Microbiology, University of Washington, Seattle, WA 98109, USA; Department of Microbiology and Immunology, University of California, San Francisco, San Francisco, CA 94158, USA; PhD Program in Biological and Biomedical Sciences, Harvard University, Boston, MA, 02115; Center for Genomic Medicine, Massachusetts General Hospital, Boston, MA, 02114; Department of Pathology, Massachusetts General Hospital, Boston, MA, 02114; Department of Pathology, Harvard Medical School, Boston, MA, 02115; Quantitative Biosciences Institute, University of California, San Francisco, San Francisco, CA 94158, USA; Innovative Genomics Institute, Berkeley, CA 94720, USA

## Abstract

CRISPR systems are prokaryotic adaptive immune systems that use RNA-guided Cas nucleases to recognize and destroy foreign genetic elements, like bacteriophages and plasmids. To overcome CRISPR immunity, phages have evolved diverse families of anti-CRISPR proteins (Acrs), each of which inhibits the nucleic acid binding or cleavage activities of specific Cas protein families. Recently, Lin et al. (2020) described the discovery and characterization of 7 different Acr families (AcrVIA1-7) that inhibit type VI-A CRISPR systems, which use the nuclease Cas13a to perform RNA-guided RNA cleavage. In this Matters Arising article, we detail several inconsistencies that question the results reported in the Lin et al. (2020) study. These include inaccurate bioinformatics analyses, as well as reported experiments involving bacterial strains that are impossible to construct. The authors were unable to provide their published strains with which we might reproduce their experiments. We independently tested the Acr sequences described in Lin et al. (2020) in two different Cas13 inhibition assays, but could not detect anti-CRISPR activity. Taken together, our data and analyses prompt us to question the claim that AcrVIA1-7 reported in Lin et al. are bona fide type VI anti-CRISPR proteins.

## Introduction

CRISPR-Cas systems provide their prokaryotic hosts with sequence-specific immunity against invading genetic elements, including bacteriophages and plasmids. To achieve immunity, short nucleotide sequences (spacers) are captured from foreign genomes and stored in the CRISPR locus, where they are transcribed and processed into small CRISPR RNAs (crRNAs) that guide the recognition and cleavage of elements matching the spacer. CRISPR systems are extremely diverse, and classified into 6 distinct types and 33 subtypes with different sequence content and mechanisms of interference (Makarova et al., 2020). Type VI CRISPR systems cleave RNA, using the RNA-guided RNase Cas13 to recognize phage mRNA transcripts.

To overcome immunity, many phages encode small anti-CRISPR (Acr) proteins, which use different mechanisms to inhibit the binding and/or enzymatic activities of Cas proteins. In June 2020, Wu and colleagues (Lin et al. 2020) reported the discovery of seven type VI-A anti-CRISPRs (AcrVIA1-AcrVIA7) encoded by strains of *Leptotrichia* and *Rhodobacter* that inhibit Cas13a from *Leptotrichia wadei* (LwaCas13a), *Leptotrichia shahii* (LshCas13a), and *Leptotrichia buccalis* (LbuCas13a). The proteins were identified bioinformatically in prophages and tested with a series of assays including *in vitro* transcription-translation (Tx-Tl), phage and plasmid targeting assays. The authors demonstrated that these Acr proteins strongly inhibited Cas13a-mediated RNA knockdown and RNA editing in human cell lines. Separately, we (A.J.M) reported that an unrelated type VI Acr (called AcrVIA1_Lse_) from a prophage of *Listeria seeligeri*, inhibits Cas13a from *Listeria seeligeri* (Meeske et al., 2020).

While reviewing the Lin et al. 2020 study, we discovered numerous troubling issues with their approach and results. Here we summarize our concerns: (i) there is no bioinformatic evidence supporting the conclusion that AcrVIA1-7 from Lin et al. are anti-CRISPR proteins, (ii) several strains reported as being used for experiments in the paper are not possible to construct due to plasmid incompatibility, (iii) none of the 7 proteins tested had anti-CRISPR activity against LwaCas13a or LbuCas13a in our own bacterial assays, and (iv) the two most potent reported proteins, AcrVIA4 and AcrVIA5, did not substantially inhibit LwaCas13a-mediated RNA knockdown in human cells.

## Results

Lin et al. used bioinformatic strategies to identify type VI Acr candidates. Specifically, they searched public sequence databases for bacterial genomes that have type VI CRISPR-Cas loci with spacers targeting another region in the same genome (often referred to as “self-targeting spacers”). A spacer and its target should not be able to stably coexist in the same cell, therefore the presence of self-targeting spacers suggests that the CRISPR system might be inhibited by an anti-CRISPR protein encoded by a prophage residing in the same genome (Rauch et al., 2017). This approach has been successfully used to identify anti-CRISPRs for DNA-targeting type II and type V CRISPR systems (Marino et al., 2018; Rauch et al., 2017; Watters et al., 2018), but not previously applied for Type VI systems, which require a transcribed target in the correct orientation for basepairing (Abudayyeh et al., 2016; Meeske et al., 2019). Lin et al. identified a single strain of *Leptotrichia wadei* (str. F0279) that possessed three self-targeting spacers associated with its type VI-A CRISPR system. Next, the authors identified three prophage regions in the *L. wadei* F0279 genome and looked for candidate anti-CRISPR genes within them. To narrow down the list of phage genes, the authors claim to have identified three anti-CRISPR-associated (*aca*) genes in the prophages (1 per prophage) and examined genes near the *aca* genes for anti-CRISPR activity. Aca proteins are so-named based on their genomic position being adjacent to known *acr* genes (Bondy-Denomy et al., 2013; Pawluk et al., 2014, 2016a, 2016b; Pinilla-Redondo et al., 2020). Mechanistically, Aca proteins are helix-turn-helix (HTH) motif containing proteins that repress *acr* expression (Birkholz et al., 2019; Osuna et al., 2020a; Stanley et al., 2019). In Fig. 1D of the original report, the authors incorrectly label the genes encoding for proteins with HTH-motifs as “*aca1, aca2*, and *aca3*” which match the names of *aca* genes previously used for *acr* discovery. Adjacent to these “*aca*” genes, the authors identified 5 putative type VI Acrs (*acrVIA1-5*) in the L. wadei F0279 genome, the most *acrs* reported in a single genome to date. The “*aca”* genes identified do encode proteins with HTH motifs, common to many phage proteins, but are not homologous to Aca1, Aca2, or Aca3, and are therefore not *aca* genes *a priori*. Thus, there is no bioinformatic basis to designate these genes *aca*s, and no basis to suggest the genes located nearby are *acr*s. The ORF depictions in Fig. 1D also mask the true size of the genes and the spacing between them (**Fig. 1**). The organization of the operons and size of the Aca proteins also does not resemble other *acr-aca* operons. For example, “*aca1”* and “*aca2*” are larger than previously identify *aca* genes, and in the case of “*aca2,”* it appears to be the first gene in an adjacent operon, and not co-encoded with the identified *acrVIA3, acrVIA4*.

**Figure 1:**
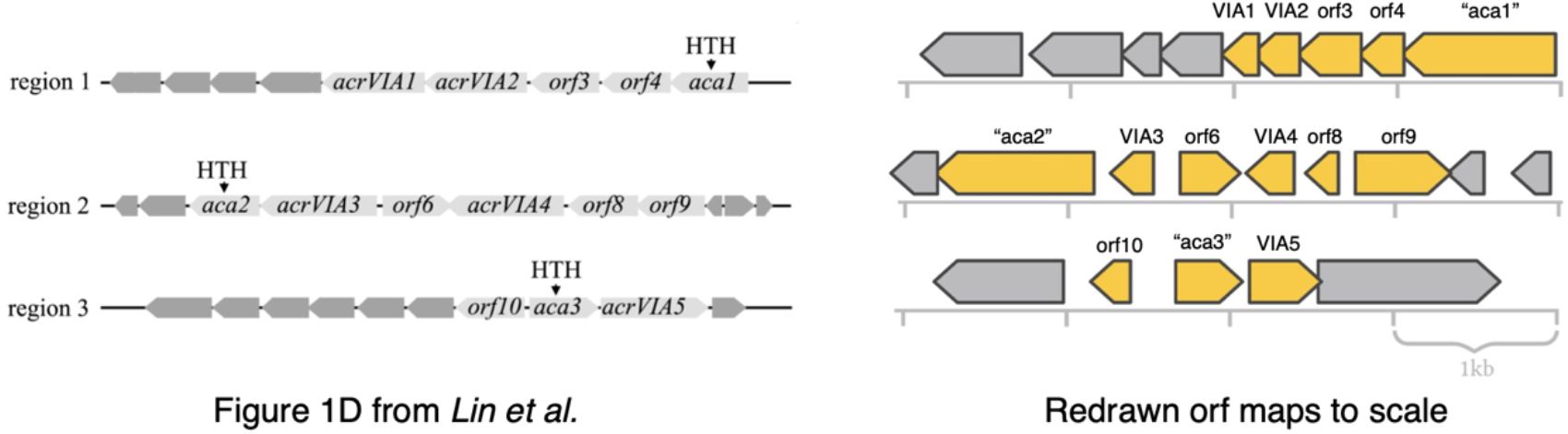
Gene organization of reported Acrs. Misnamed *aca* genes and adjacent *acr* genes as depicted in Lin et al. 2020 (left) and with ORFs redrawn to reflect their true scale and spacing (right).

Two additional type VI *acrs* (*acrVIA6, acrVIA7*) were identified by looking at genes adjacent to homologs of known *acr* genes harbored by bacterial strains also possessing type VI CRISPR loci. The discovery of *acrVIA6* was based on a nearby gene described as a homolog of the previously discovered *acrIC1* (**Fig. 2A**). However, the gene schematized *acrIC1* is misinterpreted as having 34.8% identity to the originally published *acrIC1* (Marino et al., 2018). Using BLASTp with default parameters against the nr database does not present this protein, which has few identifiable homologs. However, we were able to recreate that 34.8% identity using pairwise BLASTp that calculated 8/23 identical residues (34.78%) but to only 12% of the query protein (E-value = 0.002) (**Fig. 2B**). Similarly, the authors claimed to identify *acrIIA1* and *acrIIC4* homologs in *Leptotrichia buccalis* str. DSM1135, which led them to *acrVIA7*. Again, using the AcrIIC4 sequence (Lee et al., 2018) for BLASTp against the nr database does not present homologs in this isolate. But again we were able to recreate the erroneous identity value of 58.8% using pairwise BLASTp (10/17 residues align, over 29% of the protein) (**Fig. 2C**). We could not recreate the exact 38.9% identity of AcrIIA1 (Rauch et al., 2017) with the homolog they identify. It is likely another erroneous value generated by pairwise BLASTp.

**Figure 2:**
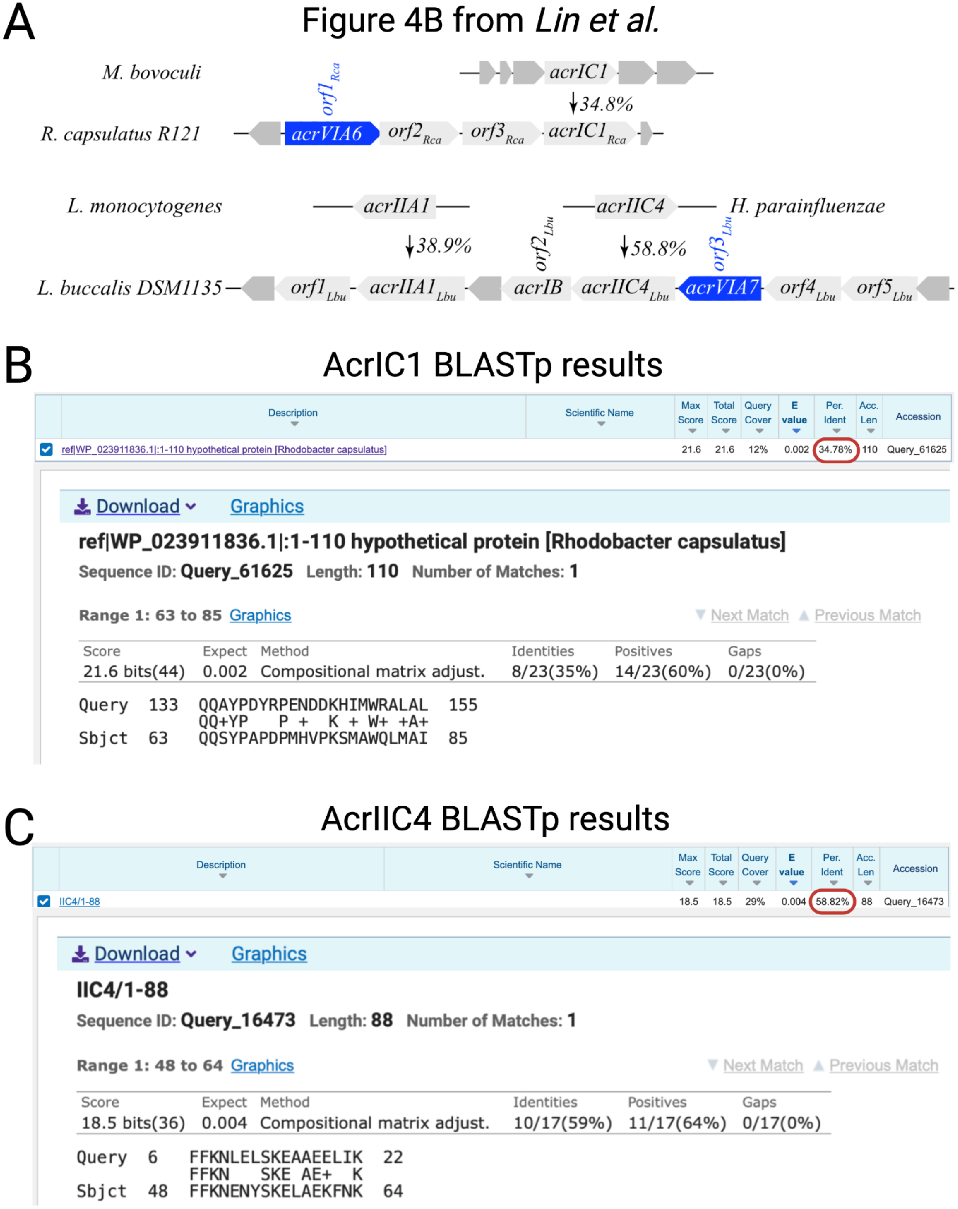
Misidentification of Acr homologs. (**A**) Reported Acr homologs with percent identity for each match indicated in Lin et al. 2020. BLASTp alignments demonstrating no significant homologs are present in the target genomes for (**B**) AcrIC1 or (**C**) AcrIIC4.

Notably, we struggled to recreate these alignments, and the authors provided no alignments of the homologs to demonstrate the percent identities. Similarly, no guidance was provided towards how BLAST searches were conducted, what the parameters were, or if PSI-BLAST was used. We conclude that every *acr* gene candidate was identified under unusual presuppositions, either through the selection of random HTH-containing proteins in prophages (*acrVIA1-5*, Fig. 1D in the original paper) and calling them *aca* genes with incorrect names (i.e., *aca1, aca2, aca3*), or through the identification of incorrect homologs of known *acr* genes (*acrVIA6-7* and *acrIB1* in Fig. 4B in the original paper).

Despite the lack of bioinformatic evidence, it remained possible that the seven candidate proteins identified by Lin et al. were indeed Acr proteins, as demonstrated through multiple experimental approaches in the original report. To test them experimentally, we first sought to reproduce some of the findings from the original study using the authors’ published reagents. All of the Acr proteins were shown by Lin et al. to reverse Cas13a-mediated interference against the *E. coli* RNA bacteriophage MS2. In hopes of repeating this experiment, we requested the published bacterial strains harboring LwaCas13a, an MS2-targeting guide, each of the individual Acrs, and an additional no-Acr control. The corresponding author did not cooperate with our request, and seven months later informed us that he could not locate the published strains. Some strains were eventually sent, however, they were not the ones we requested.

We then planned to reconstruct the strains ourselves with plasmids that were provided, but immediately realized that the strains cannot be constructed as described in the original study. MS2 phage infects the *E. coli* strain C3000, and each of the published strains contains three plasmids: (i) either an LwaCas13a or LbuCas13a expression plasmid in the vector backbone p2CT (these were generated by the Doudna lab and deposited in Addgene); (ii) an MS2-targeting guide RNA plasmid in vector backbone pCDFDuet; and (iii) an Acr expression plasmid in vector backbone pET16b. There are two major problems with these plasmid combinations. Most importantly, the p2CT backbone (East-Seletsky et al., 2016) and pET16b backbone are incompatible plasmids containing the same antibiotic resistance marker and the same origin of replication. Therefore, these strains are impossible to construct. Second, both of these backbones are designed for T7 RNA polymerase-driven gene expression for protein purification and are normally used in DE3-lysogenized host strains containing a copy of T7 RNAP (Studier et al., 1990). Strain C3000, a K12 derivative, has no copy of T7 RNAP, and therefore would drive expression of neither Cas13a nor the Acr.

Another method that Lin et al used to demonstrate Acr activity was a plasmid transformation assay in *E. coli*. Cas13a restricts the transformation of plasmids expressing target RNA, but AcrVIA1-5 were shown to relieve this restriction. Again, we encountered difficulty when trying to repeat the experiments as performed in the original study. The experiment described by Lin et al involves the use of a “self-targeting” Cas13a (LwaCas13a, LbuCas13a, or LshCas13a) plasmid, which contains a guide RNA targeting a gentamicin resistance gene on the same plasmid. The self-targeting plasmid was co-transformed into *E. coli* with or without plasmids expressing the Acrs, and transformants were only obtained in the presence of the Acr plasmids. However, such a self-targeting Cas13 plasmid should not be possible to construct in the first place, unless *cas13a* or the gentamicin resistance gene were conditionally expressed. The paper contains no information about how any of the plasmids were constructed, and there is no description of an inducible promoter used in this plasmid. We requested the strain carrying this plasmid from the corresponding author, and although it was listed among the shipped strains, we were unable to culture it in the specified antibiotics.

To test whether the seven proteins alleged to be type VI Acr proteins inhibit Cas13a, we performed plasmid transformation assays in *E. coli*. We designed plasmids carrying either LbuCas13a or LwaCas13a crRNAs targeting the kanamycin resistance marker of a second plasmid (**Fig. 3A**). To test Cas13a function, we first transformed strains harboring each Cas13a plasmid (or empty vector) with the kanR plasmid. As expected, we found that both Cas13a homologs prevented transformation with the targeted plasmid in a crRNA-dependent manner (**Fig. 3B**, no Acr control). To test the putative Acrs, we synthesized codon-optimized alleles of each gene (*acrVIA1-7*) and cloned them into the target plasmid under the control of a Ptet promoter (**Fig. 3A**). In contrast to the robust Cas13a inhibition reported by Lin et al, we found that none of the seven proteins reduced Cas13 targeting (**Fig. 3B**). As a positive control to ensure Acrs can inhibit Cas13 in this assay, we generated an LseCas13a plasmid targeting the same antibiotic resistance marker in a plasmid harboring AcrVIA1_Lse_(Meeske et al. 2020). While LseCas13a was capable of strongly interfering with a target plasmid lacking the Acr, the AcrVIA1_Lse_ target plasmid was completely resistant to LseCas13a. All of the Acrs in this experiment were expressed from the same vector backbone, using the same promoter and ribosome binding site. All of the Cas13a homologs and crRNA expression constructs were similarly matched. We therefore conclude that AcrVIA1-7 exhibit little or no anti-CRISPR activity against LbuCas13a or LwaCas13a in bacteria.

**Figure 3:**
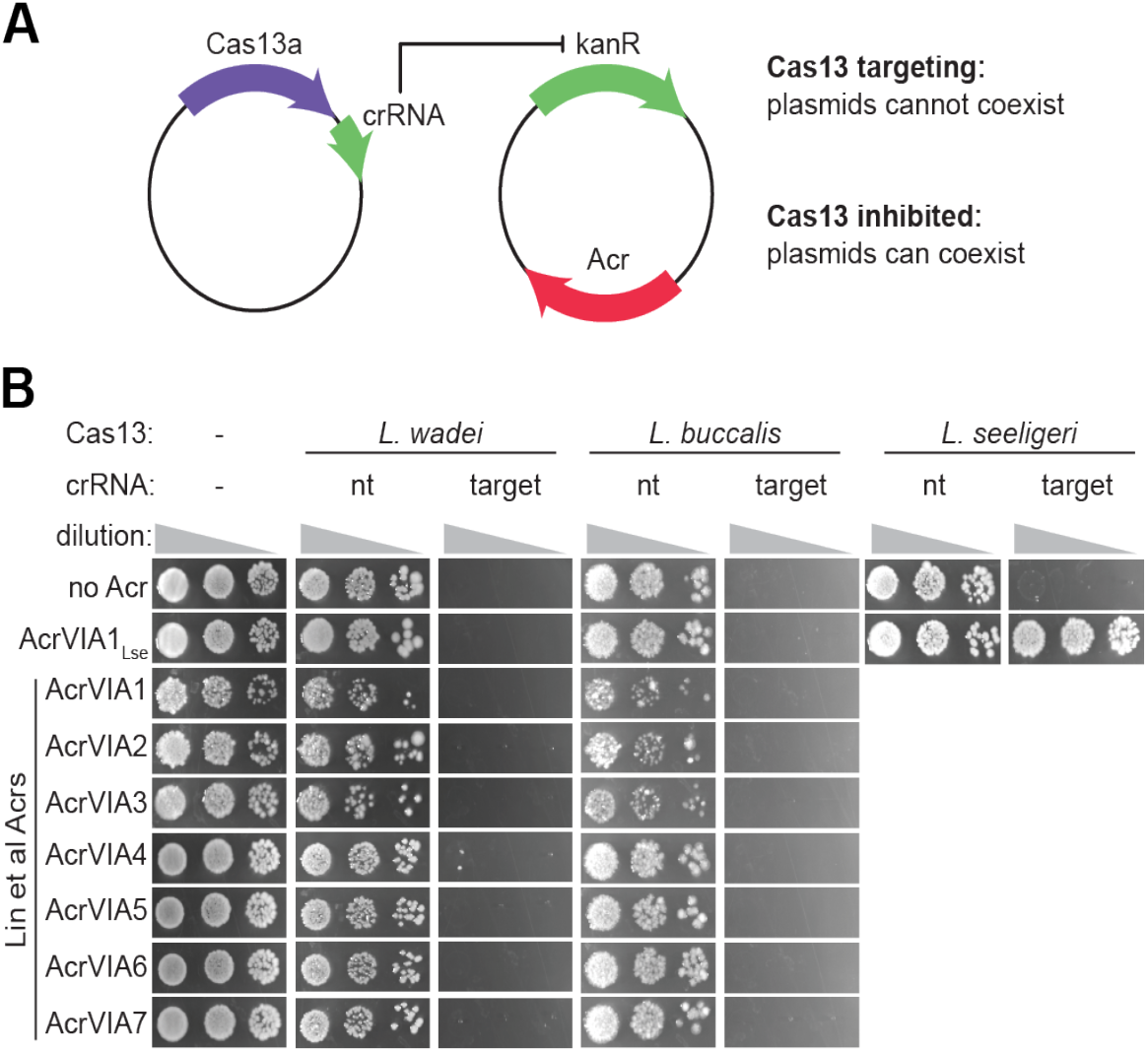
Target plasmid transformation assay to measure Acr function. (**A**) Assay schematic. *E. coli* strains carried plasmids with Cas13a from *L. wadei, L. buccalis*, or *L. seeligeri* and a crRNA targeting the kanamycin resistance marker of a second plasmid (or non-targeting crRNA control), which also candidate Acr proteins. Cas13a targeting of the kanR marker restricts transformation of the target plasmid, unless the Acr inhibits Cas13a. (**B**) Results of transformation assay. The indicated recipient strains (columns) were transformed with the indicated plasmids (rows), serially diluted and spotted onto media selecting for both plasmids. nt, non-targeting crRNA.

**Figure 4:**
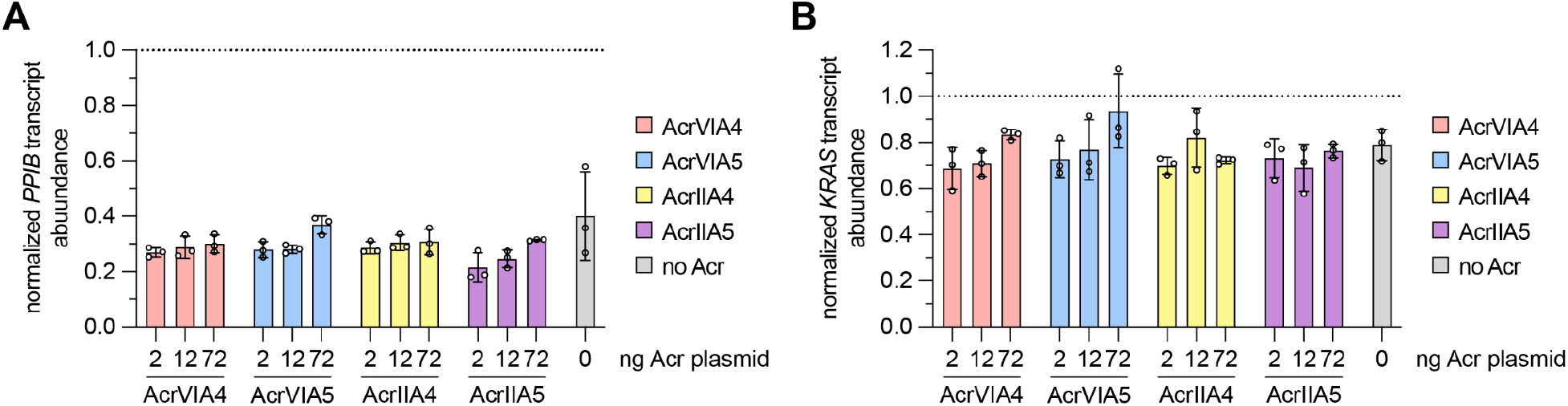
Assessment of AcrVIA-mediated inhibition of LwaCas13a. (**A, B**) Knockdown of endogenous *PPIB* (**panel A**) or *KRAS* (**panel B**) transcripts by LwaCas13a in the presence of various amounts of Acr expression plasmids was determined via RT-qPCR analysis (n = 3 biological replicates; dots represent the mean of three technical triplicate qPCR values with SD shown). *PPIB* or *KRAS* RNA levels were normalized to *ACTB* and knockdown was determined by comparison to a non-targeting LwaCas13a control. Increasing amounts of plasmids encoding the anti-CRISPR proteins AcrVIA5, AcrVIA4, AcrIIA4, and AcrIIA5 were added to a consistent amount of LwaCas13a nuclease and gRNA plasmids. The molar ratios of the Acr to LwaCas13a expression plasmids were approximately 0.12, 0.72, and 4.32 for the 2 ng, 12 ng, and 72 ng treatments, respectively.

Next, we sought to recapitulate the author’s demonstration of potent inhibition of LwaCas13a in human cells with their putative Acr proteins. Lin et al. report near-complete inhibition of LwaCas13a transcript degradation by AcrVIA5 and high levels of inhibition for several other candidate AcrVIAs. We performed experiments in HEK 293T cells with LwaCas13a and the Lin et al. crRNAs targeting endogenous *KRAS* and *PPIB* transcripts (**Figs. 4A** and **4B**, respectively), in the absence or presence of varying amounts of plasmids encoding their two most potent reported Acrs, AcrVIA5 and AcrVIA4. As negative controls, we also examined AcrIIA4 and AcrIIA5, which inhibit Cas9 (**Fig. 4**). Across the various quantities of Acr plasmids that we examined, RT-qPCR analysis at 48-hours post-transfection revealed largely inconsequential reduction of transcript degradation by AcrVIA5 or AcrVIA4; inhibition was not meaningfully different than what we observed in the presence of AcrIIA4 or AcrIIA5, which are not expected to interact with LwaCas13a. In our experiments, the highest doses of AcrVIA4 or AcrVIA5 expression plasmid were a 4.32x excess molar ratio relative to the LwaCas13a expression plasmid, which is comparable to the approximately 5.7x excess reported by Lin et al. For context, using similar construct architectures (plasmid backbones, promoters, codon usage, etc.) we previously demonstrated that AcrIIA4 and AcrIIA5 exhibit robust inhibition of SpCas9 at AcrIIA:Cas9 plasmid molar ratios as low as 0.1 (Mahendra et al., 2020; Osuna et al., 2020b). Together, our results do not support the conclusion from Lin et al. that AcrVIA5 and AcrVIA4 strongly inhibit LwaCas13a activity in human cells.

Here, we have presented multiple lines of evidence to challenge the findings put forth by Lin et al. (2020) pertaining to the discovery of seven Type VI-A Acr proteins. Bioinformatic inconsistencies are detailed, along with plasmid incompatibilities and independent experiments that could not establish the functionality of these proteins. Given the interest and utility envisioned for Acr proteins, we hope that this report is of use for the field and welcome any feedback supporting or refuting our claims.

## Acknowledgements

This work in the Bondy-Denomy lab (J.B.-D). is supported by the UCSF Program for Breakthrough Biomedical Research funded in part by the Sandler Foundation, the NIH [DP5-OD021344, R01GM127489], and DARPA HR0011-17-2-0043. M.J. is supported by the NSF GRFP Fellowship. B.P.K. acknowledges support from NIH P01-HL142494 and the Margaret Q. Landenberger Research Foundation. A.J.M was a Helen Hay Whitney postdoctoral fellow.

## Methods

### Bacterial strains and culture conditions

Plasmid culture and extraction, as well as plasmid transformation assays were performed with New England Biolabs Turbo Competent E. coli. Bacteria were grown in Lysogeny Broth (LB) supplemented with antibiotics: kanamycin (50 µg/mL), chloramphenicol (25 µg/ml), and anhydrotetracyline (100 ng/mL). 0.2% arabinose was added to induce expression of Cas13 from Para promoters.

### Plasmid construction

Cas13 plasmids for transformation assay were cloned into the vector pAM38 containing p15A origin of replication, chloramphenicol resistance marker, and an arabinose-inducible Para promoter. Coding sequences for LwaCas13a, LbuCas13a, or LseCas13a were inserted downstream of the Para promoter, and crRNA constructs were inserted downstream of the cas13 coding sequence driven by a synthetic promoter (J23119). These plasmids were constructed by three piece Gibson assembly as described below and validated by Sanger sequencing. Lists of plasmids, oligonucleotide primers, and crRNAs can be found in Supplementary Tables.

LwaCas13a plasmids: Three piece Gibson assembly using (1) SalI/HindIII-digested pAM38, (2) LwaCas13a coding sequence and Shine-Dalgarno sequence amplified from Addgene #91865 using primers oAM1496 and oAM1497, and (3) J23119-driven crRNA gBlock fragments synthesized by IDT was used to generate pAM491 (LwaCas13a kan crRNA) and pAM494 (LwaCas13a non-targeting crRNA).

LbuCas13a plasmids: Three piece Gibson assembly using (1) SalI/HindIII-digested pAM38, (2) LbuCas13a coding sequence codon-optimized for E. coli (synthesized by Genewiz) and Shine-Dalgarno sequence amplified using primers oAM211 and oAM1477, and (3) J23119-driven crRNA gBlock fragments synthesized by IDT was used to generate pAM492 (LbuCas13a kan crRNA) and pAM475 (LbuCas13a non-targeting crRNA).

LseCas13a plasmids: Three piece Gibson assembly using (1) SalI/HindIII-digested pAM38, (2) LseCas13a coding sequence codon-optimized for E. coli (synthesized by Genewiz) and Shine-Dalgarno sequence amplified using primers oAM207 and oAM994, and (3) J23119-driven crRNA gBlock fragments synthesized by IDT was used to generate pAM526 (LseCas13a kan crRNA) and pAM319 (LseCas13a non-targeting crRNA).

Acr plasmids: Acrs were expressed from pAM326 (carrying kanamycin resistance marker and pWV01 origin of replication) and driven by an anhydrotetracycline-inducible Ptet promoter. Acr coding sequences were synthesized by Genewiz, amplified using primers oAM1529 and oAM1530, and inserted with Ptet fragment into HindIII/EagI-digested pAM326 via three piece Gibson assembly to make pAM495 (AcrVIA1), pAM496 (AcrVIA2), pAM497 (AcrVIA3), pAM498 (AcrVIA4), pAM499 (AcrVIA5), pAM500 (AcrVIA6), pAM501 (AcrVIA7). The AcrVIA1Lse plasmid pAM383 was generated previously (Meeske et al. 2020).

### Plasmid transformation assay

NEB Turbo competent E. coli were transformed with either empty vector or plasmids harboring LwaCas13a, LbuCas13a, or LseCas13a and expressing non-targeting or targeting crRNAs, and selected on LB agar containing chloramphenicol and 0.2% glucose to repress Cas13a expression. Each transformed strain was made chemically competent by resuspension of exponentially growing cells in ⅕ culture volume of ice cold TFB I (10 mM CaCl2,30 mM potassium acetate, pH 5.8, 100 mM RbCl, 50mM MnCl2, 15% glycerol), incubation on ice for 15 minutes, then pelleting and resuspension in 1/25 culture volume of ice cold TFB II (10 mM MOPS pH 6.5, 10 mM RbCl, 75 mM CaCl2, 15% glycerol). Chemically competent cells were transformed by heat shock at 42°C with 100 ng Acr plasmids or empty vector control, then recovered for 1 hour in LB. Tenfold serial dilutions of recovered transformants were made and 5 µL of each dilution was spotted onto LB agar supplemented with kanamycin, chloramphenicol, arabinose, and anhydrotetracycline. Plates were photographed after 1 day of incubation at 37°C.

### Human cell culture and transfections

Human HEK 293T cells (ATCC) were cultured in Dulbecco’s Modified Eagle Medium (DMEM) supplemented with 10% heat-inactivated FBS (HI-FBS) and 1% penicillin/streptomycin. Supernatants from cell cultures were tested monthly for mycoplasma using MycoAlert PLUS (Lonza). For transfections, approximately 2 × 10^4^ cells per well were seeded in 96-well plates. Approximately 18-22 hours after seeding, transfections were performed using TransIT-X2 (Mirus) transfection reagent with 60 ng of LwaCas13a plasmid (Addgene ID 91924), 20 ng of gRNA plasmid (generated by cloning oligos into an LwaCas13a crRNA entry plasmid, LTH151, Addgene ID 171129), and with 0, 2, 12, or 72 ng of Acr plasmid. The human codon optimized Acr constructs for AcrVIA4 and AcrVIA5 were synthesized by Twist Biosciences and cloned into a pCMV backbone (plasmid IDs LTH956 and LTH957 respectively; see **Supplementary Sequences**), similar to how we previously generated the human cell expression constructs for AcrIIA4 and AcrIIA5 (Addgene IDs 133801 and 133802, respectively) (see **Supplementary Sequences**). The total amount of DNA in each transfection remained constant (152 ng), with a pCMV-null plasmid (Addgene ID 171128) used to balance DNA amounts when titrating the Acr plasmid. The plasmid mixtures were combined with 0.54 ul of TransIT-X2 and volume of Opti-MEM (ThermoFisher) to a final volume of 15 ul, incubated at room temperature for 15 minutes, and then applied to the HEK 293T cells. All experiments were performed with 3 independent biological replicates.

### RNA extraction and RT-qPCR

At 48 hours post-transfection, total RNA was extracted from transfected cells using an RNeasy Plus Mini Kit (Qiagen; cat. no. 74136). Purified RNA was reverse transcribed using a High-Capacity RNA-to-cDNA kit (ThermoFisher; cat. no. 4388950) using up to 250 ng of RNA as input. Prior to qPCR, the cDNA library for each sample was diluted 1:20. Samples for qPCR were prepared in technical triplicate with 5 ul of Fast SYBR Green Master Mix (ThermoFisher; cat. no. 4385610), 3 ul diluted cDNA, and 2 ul qPCR primer pairs specific to the target (IDT; **Supplementary Table 1**). Control reactions amplifying ACTB were set up in parallel for each cDNA library. Reaction cycling was performed using a Roche LightCycler480. To determine the fold change of each sample, the expression levels were normalized to a negative control transfection containing LwaCas13a plasmid and an empty pUC19 backbone plasmid (Addgene ID 133961). Each qPCR reaction was analyzed for purity by melting curve analysis to confirm a single PCR product.

**Supplementary Table 1.**
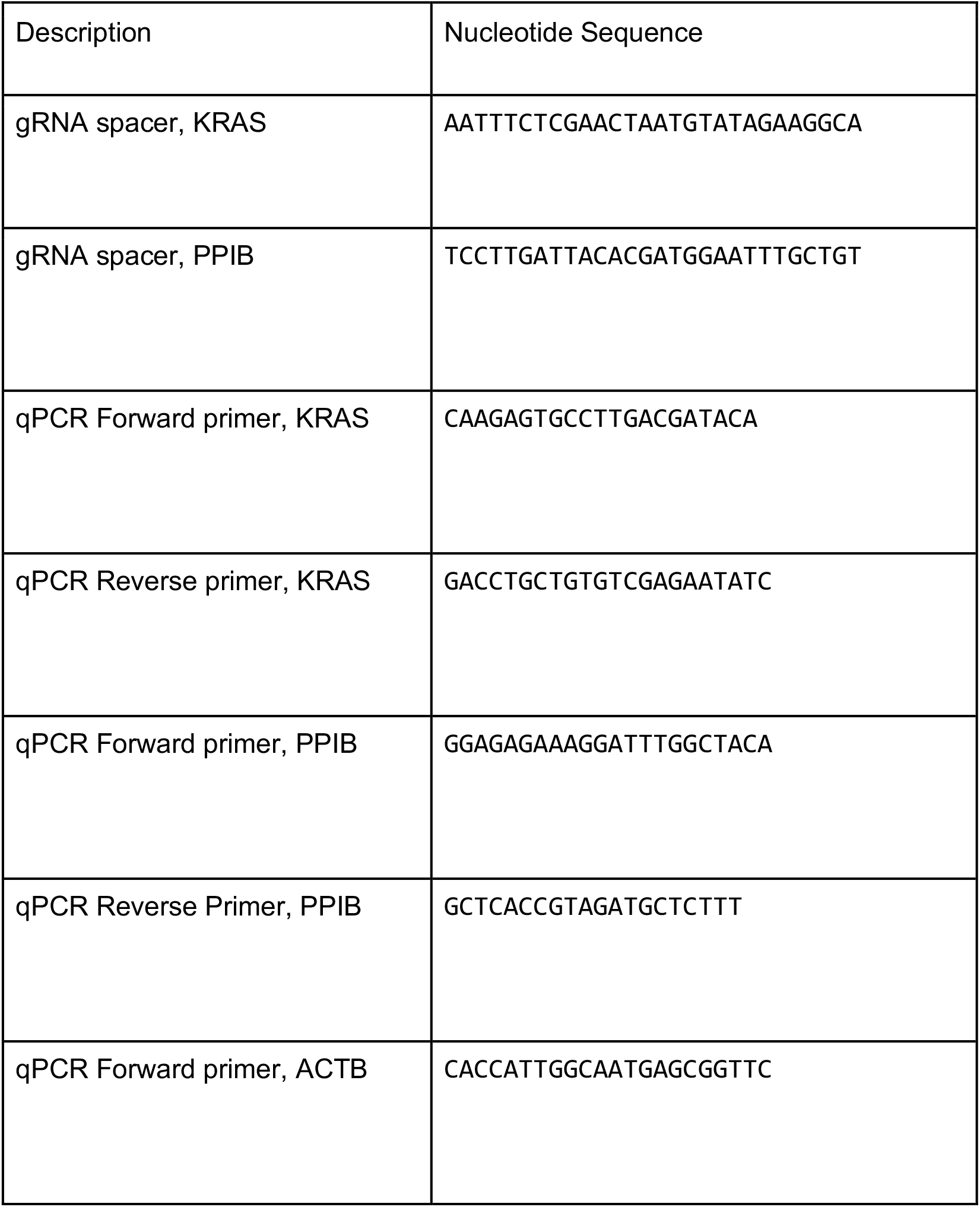
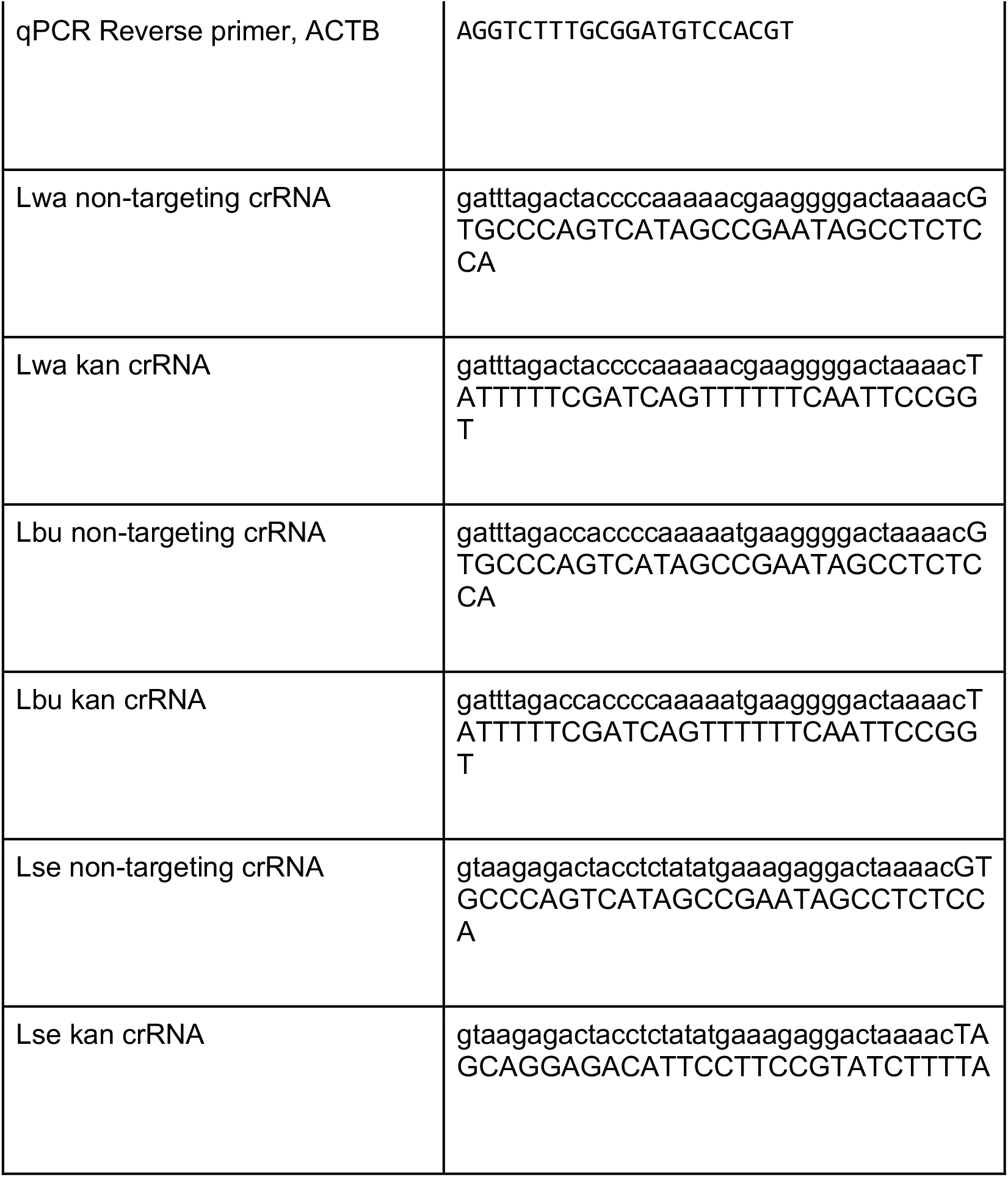
Sequences of gRNAs and oligonucleotides.

## Supplementary Sequences

Amino acid and nucleotide sequences for constructs used in human cell experiments. Linker in red. NLS in green.

>pCMV-T7-AcrVIA4 (LTH956)

MDKANRCLKAKDKILNILEKEEITLDEFNNISKDIAKEYVEKAVLKPKDIAERIINMVKNAKSISFDE LASEISEE

ATGGACAAGGCCAACAGATGCCTGAAGGCCAAGGACAAGATCCTGAACATCCTGGAGAAG GAGGAGATCACCCTGGACGAGTTCAACAACATCAGCAAGGACATCGCCAAGGAGTACGTG GAGAAGGCCGTGCTGAAGCCCAAGGACATCGCCGAGAGAATCATCAACATGGTGAAGAAC GCCAAGAGCATCAGCTTCGACGAGCTGGCCAGCGAGATCAGCGAGGAG

>pCMV-T7-AcrVIA5 (LTH957)

MERNFKKVTENTGRKEVFKVMHDKVEIINDFNTNEKREARIIFHDQKIYVILYQNLNFEELKWLN FYILIYGNQSYGKNTFFEFKLNKNNLIYHLQVWNIIENKKFKSKSISLLVKALSSKAGV

ATGGAGAGAAACTTCAAGAAGGTGACCGAGAACACCGGCAGAAAGGAGGTGTTCAAGGTG ATGCACGACAAGGTGGAGATCATCAACGACTTCAACACCAACGAGAAGAGAGAGGCCAGA ATCATCTTCCACGACCAGAAGATCTACGTGATCCTGTACCAGAACCTGAACTTCGAGGAGC TGAAGTGGCTGAACTTCTACATCCTGATCTACGGCAACCAGAGCTACGGCAAGAACACCTT CTTCGAGTTCAAGCTGAACAAGAACAACCTGATCTACCACCTGCAGGTGTGGAACATCATC GAGAACAAGAAGTTCAAGAGCAAGAGCATCAGCCTGCTGGTGAAGGCCCTGAGCAGCAAG GCCGGCGTG

>pCMV-T7-AcrIIA4-NLS(SV40) (KAC200; Addgene ID 133801)

MGNINDLIREIKNKDYTVKLSGTDSNSITQLIIRVNNDGNEYVISESENESIVEKFISAFKNGWNQ EYEDEEEFYNDMQTITLKSELNGSGGGGSGPKKKRKV

ATGGGCAACATCAACGACCTGATCAGAGAGATCAAGAACAAGGACTACACCGTGAAGCTG AGCGGCACCGACAGCAACAGCATCACCCAGCTGATCATCAGAGTGAACAACGACGGCAAC GAGTACGTGATCAGCGAGAGCGAGAACGAGAGCATCGTGGAGAAGTTCATCAGCGCCTTC AAGAACGGCTGGAACCAGGAGTACGAGGACGAGGAGGAGTTCTACAACGACATGCAGAC CATCACCCTGAAGAGCGAGCTGAACGGATCCGGCGGTGGAGGCAGTGGGCCCAAGAAGA AGAGGAAAGTC

>pCMV-T7-AcrIIA5-NLS(SV40) (KAC203; Addgene ID 133802)

MAYGKSRYNSYRKRNFSISDNQRREYAKKMKELEQAFENLDGWYLSSMKDSAYKDFGKYEIR LSNHSADNRYHDLENGRLIVNVKASKLNFVDIIENKLGKIIEKIDTLDLDKYRFINATKLERDIKCYY KGYKTKKDVIGSGGGGSGPKKKRKV

ATGGCCTACGGCAAGAGCAGATACAACAGCTACAGAAAGAGAAACTTCAGCATCAGCGAC AACCAGAGAAGAGAGTACGCCAAGAAGATGAAGGAGCTGGAGCAGGCCTTCGAGAACCT GGACGGCTGGTACCTGAGCAGCATGAAGGACAGCGCCTACAAGGACTTCGGCAAGTACG AGATCAGACTGAGCAACCACAGCGCCGACAACAGATACCACGACCTGGAGAACGGCAGAC TGATCGTGAACGTGAAGGCCAGCAAGCTGAACTTCGTGGACATCATCGAGAACAAGCTGG GCAAGATCATCGAGAAGATCGACACCCTGGACCTGGACAAGTACAGATTCATCAACGCCA CCAAGCTGGAGAGAGACATCAAGTGCTACTACAAGGGCTACAAGACCAAGAAGGACGTGA TCGGATCCGGCGGTGGAGGCAGTGGGCCCAAGAAGAAGAGGAAAGTC

>pC034-LwCas13a-msfGFP-2A-Blast was a gift from Feng Zhang (Addgene Plasmid #91924)

## References

Abudayyeh, O.O., Gootenberg, J.S., Konermann, S., Joung, J., Slaymaker, I.M., Cox, D.B.T., Shmakov, S., Makarova, K.S., Semenova, E., Minakhin, L., et al. (2016). C2c2 is a single-component programmable RNA-guided RNA-targeting CRISPR effector. Science 353, aaf5573.

Birkholz, N., Fagerlund, R.D., Smith, L.M., Jackson, S.A., and Fineran, P.C. (2019). The autoregulator Aca2 mediates anti-CRISPR repression. Nucleic Acids Res. 47, 9658–9665.

Bondy-Denomy, J., Pawluk, A., Maxwell, K.L., and Davidson, A.R. (2013). Bacteriophage genes that inactivate the CRISPR/Cas bacterial immune system. Nature 493, 429–432.

East-Seletsky, A., O’Connell, M.R., Knight, S.C., Burstein, D., Cate, J.H.D., Tjian, R., and Doudna, J.A. (2016). Two distinct RNase activities of CRISPR-C2c2 enable guide-RNA processing and RNA detection. Nature 538, 270–273.

Lee, J., Mir, A., Edraki, A., Garcia, B., Amrani, N., Lou, H.E., Gainetdinov, I., Pawluk, A., Ibraheim, R., Gao, X.D., et al. (2018). Potent Cas9 Inhibition in Bacterial and Human Cells by AcrIIC4 and AcrIIC5 Anti-CRISPR Proteins. MBio 9.

Lin, P., Qin, S., Pu, Q., Wang, Z., Wu, Q., Gao, P., Schettler, J., Guo, K., Li, R., Li, G., et al. (2020). CRISPR-Cas13 Inhibitors Block RNA Editing in Bacteria and Mammalian Cells. Mol. Cell 78, 850–861.e5.

Mahendra, C., Christie, K.A., Osuna, B.A., Pinilla-Redondo, R., Kleinstiver, B.P., and Bondy-Denomy, J. (2020). Broad-spectrum anti-CRISPR proteins facilitate horizontal gene transfer. Nat Microbiol 5, 620–629.

Makarova, K.S., Wolf, Y.I., Iranzo, J., Shmakov, S.A., Alkhnbashi, O.S., Brouns, S.J.J., Charpentier, E., Cheng, D., Haft, D.H., Horvath, P., et al. (2020). Evolutionary classification of CRISPR-Cas systems: a burst of class 2 and derived variants. Nat. Rev. Microbiol. 18, 67–83.

Marino, N.D., Zhang, J.Y., Borges, A.L., Sousa, A.A., Leon, L.M., Rauch, B.J., Walton, R.T., Berry, J.D., Joung, J.K., Kleinstiver, B.P., et al. (2018). Discovery of widespread type I and type V CRISPR-Cas inhibitors. Science 362, 240–242.

Meeske, A.J., Nakandakari-Higa, S., and Marraffini, L.A. (2019). Cas13-induced cellular dormancy prevents the rise of CRISPR-resistant bacteriophage. Nature 570, 241–245.

Meeske, A.J., Jia, N., Cassel, A.K., Kozlova, A., Liao, J., Wiedmann, M., Patel, D.J., and Marraffini, L.A. (2020). A phage-encoded anti-CRISPR enables complete evasion of type VI-A CRISPR-Cas immunity. Science.

Osuna, B.A., Karambelkar, S., Mahendra, C., Sarbach, A., Johnson, M.C., Kilcher, S., and Bondy-Denomy, J. (2020a). Critical Anti-CRISPR Locus Repression by a Bi-functional Cas9 Inhibitor. Cell Host Microbe 28, 23–30.e5.

Osuna, B.A., Karambelkar, S., Mahendra, C., Christie, K.A., Garcia, B., Davidson, A.R., Kleinstiver, B.P., Kilcher, S., and Bondy-Denomy, J. (2020b). Listeria Phages Induce Cas9 Degradation to Protect Lysogenic Genomes. Cell Host Microbe 28, 31–40.e9.

Pawluk, A., Bondy-Denomy, J., Cheung, V.H.W., Maxwell, K.L., and Davidson, A.R. (2014). A new group of phage anti-CRISPR genes inhibits the type I-E CRISPR-Cas system of Pseudomonas aeruginosa. MBio 5, e00896.

Pawluk, A., Staals, R.H.J., Taylor, C., Watson, B.N.J., Saha, S., Fineran, P.C., Maxwell, K.L., and Davidson, A.R. (2016a). Inactivation of CRISPR-Cas systems by anti-CRISPR proteins in diverse bacterial species. Nature Microbiology 1, 1–6.

Pawluk, A., Amrani, N., Zhang, Y., Garcia, B., Hidalgo-Reyes, Y., Lee, J., Edraki, A., Shah, M., Sontheimer, E.J., Maxwell, K.L., et al. (2016b). Naturally Occurring Off-Switches for CRISPR-Cas9. Cell 167, 1829–1838.e9.

Pinilla-Redondo, R., Shehreen, S., Marino, N.D., Fagerlund, R.D., Brown, C.M., Sørensen, S.J., Fineran, P.C., and Bondy-Denomy, J. (2020). Discovery of multiple anti-CRISPRs highlights anti-defense gene clustering in mobile genetic elements. Nat. Commun. 11, 5652.

Rauch, B.J., Silvis, M.R., Hultquist, J.F., Waters, C.S., McGregor, M.J., Krogan, N.J., and Bondy-Denomy, J. (2017). Inhibition of CRISPR-Cas9 with Bacteriophage Proteins. Cell 168, 150–158.e10.

Stanley, S.Y., Borges, A.L., Chen, K.-H., Swaney, D.L., Krogan, N.J., Bondy-Denomy, J., and Studier, F.W., Rosenberg, A.H., Dunn, J.J., Dubendorff, J.W. (1990). Use of T7 RNA polymerase to direct expression of cloned genes. Methods Enzymol. 185:60–89

Davidson, A.R. (2019). Anti-CRISPR-Associated Proteins Are Crucial Repressors of Anti-CRISPR Transcription. Cell 178, 1452–1464.e13.

Watters, K.E., Fellmann, C., Bai, H.B., Ren, S.M., and Doudna, J.A. (2018). Systematic discovery of natural CRISPR-Cas12a inhibitors. Science 362, 236–239.

